# Multi-omics data and analysis reveal the formation of key pathways of different colors in *Torenia fournieri* flowers

**DOI:** 10.1101/2023.06.19.545640

**Authors:** Jiaxing Song, Haiming Kong, Jing Yang, Jiaxian Jing, Siyu Li, Nan Ma, Rongchen Yang, Yuman Cao, Yafang Wang, Tianming Hu, Peizhi Yang

**Affiliations:** College of Grassland Agriculture, Northwest A&F University, Yangling, Shaanxi, 712100, China

**Author notes:** **Email addresses:**.

**Keywords:** *Torenia fournieri*, Genome, RNA-seq;Flavonoids, Anthocyanins, Flower color

## Abstract

*Torenia fournieri* Lind. is an ornamental plant, popular for its numerous flowers and variety of colors. However, its genomic evolution, as well as the genetic and metabolic basis of flower color formation, remain poorly understood. Here we report a chromosome-level reference genome of *T. fournieri* comprising 164.4 Mb. Phylogenetic analysis revealed the phylogenetic placement of the species, and comparative genomics analysis indicated that *T. fournieri* shared a whole genome duplication (WGD) event with *Antirrhinum majus*. Through joint transcriptomics and metabolomics analyses, we characterized the differential genes and metabolites in the anthocyanin synthesis pathway in five *T. fournieri* varieties. We identified many metabolites related to pelargonidin, peonidin, and naringenin in Rose (R) color samples. On the other hand, the blue (B) and blue-violet (D) color samples contained many metabolites related to petunidin, cyanidin, quercetin, and malvidin. The formation of different flower colors in *T. fournieri* involves multiple genes and metabolites. We analyzed the results and obtained significantly different genes and metabolites related to the biosynthesis of flavonoids and anthocyanins, which are key metabolites in the formation of different flower colors. Our *T. fournieri* genome data provide a basis for studying the differentiation of this species and provide a valuable model genome enabling genetic studies and genomics-assisted breeding of *T. fournieri*.

**Highlight:** The genome of *Torenia fournieri* was reported for the first time, and the formation mechanism of different colors in *Torenia fournieri* flowers was analyzed by genomics, transcriptomics and metabolomics.

## Introduction

*Torenia fournieri* Linden. *ex* Fourn. (also known as Wishbone flower) is an annual herb of the family Linderniaceae, suitable for warm and humid climates, grown mainly in tropical and subtropical regions(Chen *et al*., 2021; Nishihara *et al*., 2013). *T. fournieri* is a popular ornamental plant that comes in a wide variety of colors, from white and yellow to blue, violet, and lavender(Guan *et al*., 2021)3]. *T. fournieri* is also an experimental model plant(Aida, 2008). The semi-naked embryo sac structure of *T. fournieri* is conducive to the separation of egg cells and reduces the technical barriers *in vitro* fertilization operations, serving as a model plant in angiosperm flower organ development and fertilization biology research(Aida, 2008; Higashiyama *et al*., 2006; Higashiyama *et al*., 1998). In horticultural plants, flower traits, such as petal color and shape, are considered to be very important for their commercial value(Nishihara *et al*., 2013). Flower color is one of the key traits for *T. fournieri* genetic improvement to further increase the commercialization of its cultivars. Currently, no *T. fournieri* reference genome sequence *T. fournieri* has been published, which hinders its molecular design breeding.

In recent years, much effort has been placed into understanding the molecular and biochemical mechanisms of pigment formation in *T. fournieri* flowers. Flavonoids are the main compounds responsible for the color of most plants. The genes involved in the flavonoid biosynthesis pathway play a crucial role in regulating plant color(Iwashina, 2015). The dihydroflavonol-4-reductase (DFR) is an enzyme in the flavonoid biosynthesis pathway with key roles in regulating flower color(Tian *et al*., 2017). It was reported that DFR gene inactivation *T. fournieri* resulted in flavonoid accumulation, resulting in a deeper blue flower color (Aida *et al*., 2000b). Chalcone synthase (CHS) is the first enzyme to act on the flavonoid pathway and is key for the biosynthesis of precursors to other flavonoids (Liu *et al*., 2021). *TfCHS* gene was overexpressed in *T. fournieri* by transgenic technology to alter the changes in its flower color, and obtained with new characters in flower color(Aida *et al*., 2000a; Suzuki *et al*., 2000). Flavonoid 3-hydroxylase (F3H) is a key enzyme for anthocyanin synthesis in *T. fournieri* flowers(Nishihara *et al*., 2014). The absence of *TfF3H* led to reduced petal anthocyanin levels and resulted in a white petal color. Overexpression of the *F3H* gene in Crown White (CrW, white-flowered cultivar of *T. fournieri*) resulted in pink petals, a color arising from pelargonidin derivatives that lack B-ring hydroxylation(Nishihara *et al*., 2014). In the entire anthocyanin biosynthesis pathway, anthocyanin synthase (ANS) catalyzes the final step of color formation, involving the conversion of colorless anthocyanins into colored anthocyanins(Shi *et al*., 2015). The *ANS* gene expression was reduced by RNAi technology in summer *T. fournieri*, resulting in a white flower color(Nakamura *et al*., 2006). These transgenic functional studies have contributed to our understanding of the gene functions involved in *T. fournieri* flower color formation. However, as the complete reference genome of *T. fournieri* has not yet been published, it hinders the further study of the gene regulatory mechanism controlling flower color in *T. fournieri*. Therefore, assembling the reference genome of *T. fournieri* could provide the basis for the establishment of genetic engineering and genomics-assisted breeding and improve genotype to phenotype association studies.

Here, we obtained a chromosome-level assembly of the *T. fournieri* genome by combining Illumina, PacBio, and Hi-C sequencing assembly. In addition, we performed a relatively complete annotation using the assembled genome, constructed a phylogenetic tree including the main species of Plantaginaceae, Linderniaceae, and Labiatae, and assessed the evolutionary relationship between *T. fournieri* and whole-genome duplication (WGD) events. We used comparative genomics to determine the phylogenetic position of *T. fournieri*, which shared WGDs with *A. majus*. Through a multi-omics analysis combining genomics, transcriptomics, and metabolomics, we analyzed the differences in the flavonoid and anthocyanin metabolic pathways in different flower-colored genotypes. We obtained differential genes and metabolites related to the color formation of *T. fournieri*. The results of this study provide a valuable genomic basis for molecular genetic studies and the breeding of *T. fournieri*.

## Materials and methods

### Plant materials and genomic sequencing

The plant materials used in this study were grown in the greenhouse of the college of Grassland Agriculture, Northwest A&F University. The DNAsecure Plant Kit (TIANGEN) was used to extract DNA from 7-week-old *T. fournieri* fresh leaves. DNA samples of sufficient quality were prepared by a Covaris sonicator to complete the library preparation. Next-generation sequencing (NGS) was performed using the Illumina NovaSeq 6000 platform. Furthermore, we obtained high-quality single molecular sequencing reads through the PacBio Sequel platform. After the Hi-C library was constructed according to standard procedures, it was sequenced on an Illumina NovaSeq 6000 sequencer. We used Jellyfish (2.1.4) to generate the 21-mer count distribution of NGS reads(Marçais and Kingsford, 2011) and then estimated the genome size, heterozygosity, and repeat content according to the analysis model provided by GenomeScope(Ranallo-Benavidez *et al*., 2020). The PacBio reads were corrected using the falcon software(Chin *et al*., 2016), and were then assembled to obtain the genome sequence. This sequence was then used for a second round of error corrections using Pilon(Walker *et al*., 2014). We used BWA (0.7.10-r789) to align the Hi-C sequencing paired-end reads with the contigs of the assembled genome(Li and Durbin, 2010). We used the LACHESIS software to group, rank, and orient the genomic contigs sequences(Burton *et al*., 2013). To evaluate the accuracy, continuity, connectivity, and completeness of the *T. fournieri* genome assembly results, we used BUSCO software (Simão *et al*., 2015).

### Gene prediction and function annotation

The Maker software(Cantarel *et al*., 2008) was used to annotate the *T. fournieri* genome^7^, and AUGUSTUS 3.3 was used for de novo gene prediction, and the complete annotation information was obtained(Stanke *et al*., 2006). To identify transposable elements, we used the RepeatMasker(Tarailo-Graovac and Chen, 2009) and RepeatModeler(Flynn *et al*., 2020) for the identification and classification of transposable elements (TEs) sequences in the *T. fournieri* genome. The BLASTN was used to map the *A. thaliana* protein sequences into the *T. fournieri* genome and then used GENEWISE 2.4.1 to predict accurate gene models(Li *et al*., 2015). Gene function annotation mainly included two steps: sequence similarity-based functional annotation information and HMM model-based protein domain annotation information. The diamond software(Buchfink *et al*., 2015) was used to compare the genes and proteins in the *T. fournieri* genome against databases such as the NCBI non-redundant protein sequence (Nr), SwissProt(Bairoch and Apweiler, 2000), Gene Ontology (GO)(Ashburner *et al*., 2000), Kyoto Encyclopedia of Genes and Genomes (KEGG)(Qiu, 2013), KOG and Pfam(Nawrocki *et al*., 2014).

### Comparative genome analysis between species and WGD analysis

We downloaded the genome data of 11 species from Phytozome (https://phytozome.jgi.doe.gov/pz/portal.html) and selected *Amborella trichopoda* and *Vitis vinifera* as the outgroups of *T. fournieri* for comparative genome analyses. Orthofinder was run with default settings to identify homologous genes among the 12 species(Emms and Kelly, 2019). According to the Orthofinder analysis results, the jvenn software was used to map the homologous genes in *S. bowleyana*, *S. cusia*, *O. majorana*, *L. philippensis*, *S. baicalensis* and *T. fournieri*(Bardou *et al*., 2014). We used the mcmctree tool in the PAML software package to construct the 12-species phylogenetic tree together with fossil time-calibrated phylogenetic trees, calibrated with the angiosperm *A. trichopoda* (∼179.0 - 199.1 MYA) and the labiata *S. baicalensis-O. basilicum*(∼31.6 - 73.1 MYA)(Yang, 2007). The assessment of gene family expansion and contraction was performed using CAFE v5 (default settings)(Mendes *et al*., 2020), and was based on gene family clustering statistics and species phylogenetic trees at divergence time. Finally, an evolutionary tree was constructed using the online iTOL software (Interactive Tree Of Life)(Letunic and Bork, 2016).

To obtained orthologous gene pairs using the WGDI software(Sun *et al*., 2022) and calculated the synonymous substitution rate (Ks) for each synonymous gene pair, according to the gene family phylogeny using the KaKs Calculator software(Wang *et al*., 2010). Density maps of the Ks values distribution across species were plotted using the ggplot2 package for R to identify whole-genome duplication events (WGDs). Genome-wide blocks of collinearity within *T. fournieri* were identified using MCScan(Wang *et al*., 2012). Genome collinearity was finally visualized by the Python version of MCScan (Python version).To analyze retrotransposons with long terminal repeats (LTR), we used LTR_finder(Ou and Jiang, 2017) and the LTRharvest software (Ellinghaus *et al*., 2008).

### Transcription analysis

We collected *T. fournieri* flowers of five varietal colors and performed transcriptome sequencing analysis. The five more common colors of the *T. fournieri* flowers are white (marked as W, the same below), Rose (R), lemon drop (Y), blue and white (B), and deep blue (D). Total RNA was extracted from corollas of the different flowers by the Trizol method(Rio *et al*., 2010), and the library was constructed and sequenced using the Illumina platform. The fastp software was uesd to perform quality control on raw reads to obtain Clean Reads(Chen *et al*., 2018), and used HISAT to align the Clean Reads with the *T. fournieri* genome to obtain position information on the reference genome or gene(Kim *et al*., 2015). StringTie(Shumate *et al*., 2022) was used to assemble reads into transcripts, GffCompare(Pertea and Pertea, 2020) was used to compare with the genome annotation information, and finally, new transcripts or new genes were obtained. The diamond(Buchfink *et al*., 2021) software was used to align all genes with the KEGG, GO, NR, Swiss-Prot, TrEMBL, and KOG database sequences to obtain annotation results, and the alignment cutoff was an E-value of 1e-5. The featurecounts v1.6.2 was used to calculate gene alignment and FPKM(Liao *et al*., 2013). Differential expression between the two groups was analyzed using DESeq2(Love *et al*., 2014), and P-values were corrected using the method of Benjamini & Hochberg(Love *et al*., 2014). The |log2foldchange| >1 was used as the threshold for the DEGs.

### Analysis of the cytochromeP450 and R2R3-MYB gene families

The Hidden Markov Model (HMM) containing the p450 (PF00067) and MYB (PF00249) domains was obtained from the Pfam database. The domains were aligned with the HMMER software(Eddy and Eddy, 2015). We downloaded the sequences of the Arabidopsis P450 and R2R3-MYB proteins from the *A. thaliana* database (https://www.arabidopsis.org/index.jsp) and queried these sequences against the protein sequences of *T. fournieri* using BlastP software(Boratyn *et al*., 2013) (E-value≤1e-5). The obtained alignment results were combined and deduplicated, and the obtained protein sequences were screened. The results were compared by the Muscle software(Edgar, 2004), and an evolutionary tree was constructed using the MFP mode of the iqtree software (UFBoot is 1000)(Nguyen *et al*., 2014). Based on the taxonomic information of the *A. thaliana* P450 and R2R3-MYB subfamilies, the taxonomic information of the respective subfamilies in *T. fournieri* was determined, and they were named according to theirc chromosome positions. The tandem repeats of the P450 and R2R3-MYB gene family sequences in *T. fournieri* and *A. thaliana* were analyzed using the MCScanX software(Wang *et al*., 2012).

### Metabolites analysis

We selected *T. fournieri* flowers of five different colors to be freeze-dried in a vacuum freeze dryer (Scientz-100F). The samples were pulverized with a mixing mill, dissolved in a methanol solution, and centrifuged. The extracted supernatant was filtered (SCAA-104, pore size 0.22 μm) before UPLC-MS/MS analysis. Flavonoid and anthocyanin metabolite contents were detected by MetWare (http://www.metware.cn/) based on the AB Sciex QTRAP 6500 UPLC-MS/MS platform. Mass spectral data were processed using the Analyst 1.6.3 software. Comparative analysis of the two groups in the VIP (VIP ≥ 1) and absolute Log2FC (|Log2FC| ≥ 1.0) to determined differential metabolites. VIP values were extracted from the OPLS-DA result and were generated using the R package MetaboAnalystR(Chong *et al*., 2019).

### Transcriptome and metabolome conjoint analysis

Combined with the metabolome and transcriptome analysis results, the DEGs and differential metabolites of the same group of samples were co-mapped to the corresponding KEGG pathways. The main pathways mapped to KEGG_map were Flavonoid biosynthesis (ko00941), Anthocyanin biosynthesis (ko00942), and Flavone and flavonol biosynthesis (ko00944). To evaluate the differential genes and differential metabolite correlations, we used the cor function in R to calculate the Pearson correlation coefficients of genes and metabolites. The criterion of the results was correlation coefficients > 0.80 and a p-value < 0.05.

### RT-qPCR Analysis

First-strand cDNA synthesis was performed with the FastPure Plant Total RNA Isolation Kit (suitable for polysaccharide & polyphenolic rich tissues). The total RNA extracted was also used for RNA-seq library construction. Gene-specific primers were designed using Primer Premier 5.0 (Table S23). Real-time qPCR was performed using the Roche LightCycler 480II Real-Time PCR System (Roche, Basel, Switzerland) with the SYBR Green PCR Master Mix. Relative transcript levels were calculated according to the 2^−ΔΔCt^ method(Livak and Schmittgen, 2001).

## Results

### Genome sequencing and assembly

We generated 7.2 Gb Illumina 150 bp pairedend reads data(Table S1). The genome size was estimated to be approximately 187.0 Mb, using the software GenomeScope based on the kmer method (k = 21), with a heterozygosity rate of 0.81% (Fig. S2). Moreover, a total of 2.2 million reads larger than 500 bp were obtained by PacBio Sequel sequencing, with a coverage depth of approximately 79 X (Fig. S3). 149,029 reads (about 52% of the total) were larger than 5 kb in length, of which 81.57% had an average base length of 10 kb (Fig. S3). The Falcon software was used to assemble the PacBio sequencing reads, and the Pilon software was used for further genome polishing using the Illumina reads data. Finally, we obtained a genome size of 164.4Mb with a Contig N50 of 918.3kb (Table S2).

The Hic data were aligned with the assembled genome sequence using BWA (Tables S3), and divided into 9 chromosomes using LACHESIS software (Fig. S4). After Hi-C linkage data analysis, a total of 158.29 Mb sequence length was assigned to chromosomes, accounting for 96.32% of the total sequence length (Table S4). The longest chromosome was 22.1Mb, and the shortest was 13.9Mb (Fig. 1; Table S4). To evaluate the assembled genome quality, 91.64% of the sequences obtained by the BUSCO software were fully present in the *T. fournieri* genome (total number of orthologous genes in the GenBank 1614), while 5.08% and 3.28% of the BUSCO genes were partially present or absent, respectively (Fig. S5; Table S5). The above results strongly support the reliability and integrity of the *T. fournieri* genome assembly.

**Fig. 1.**
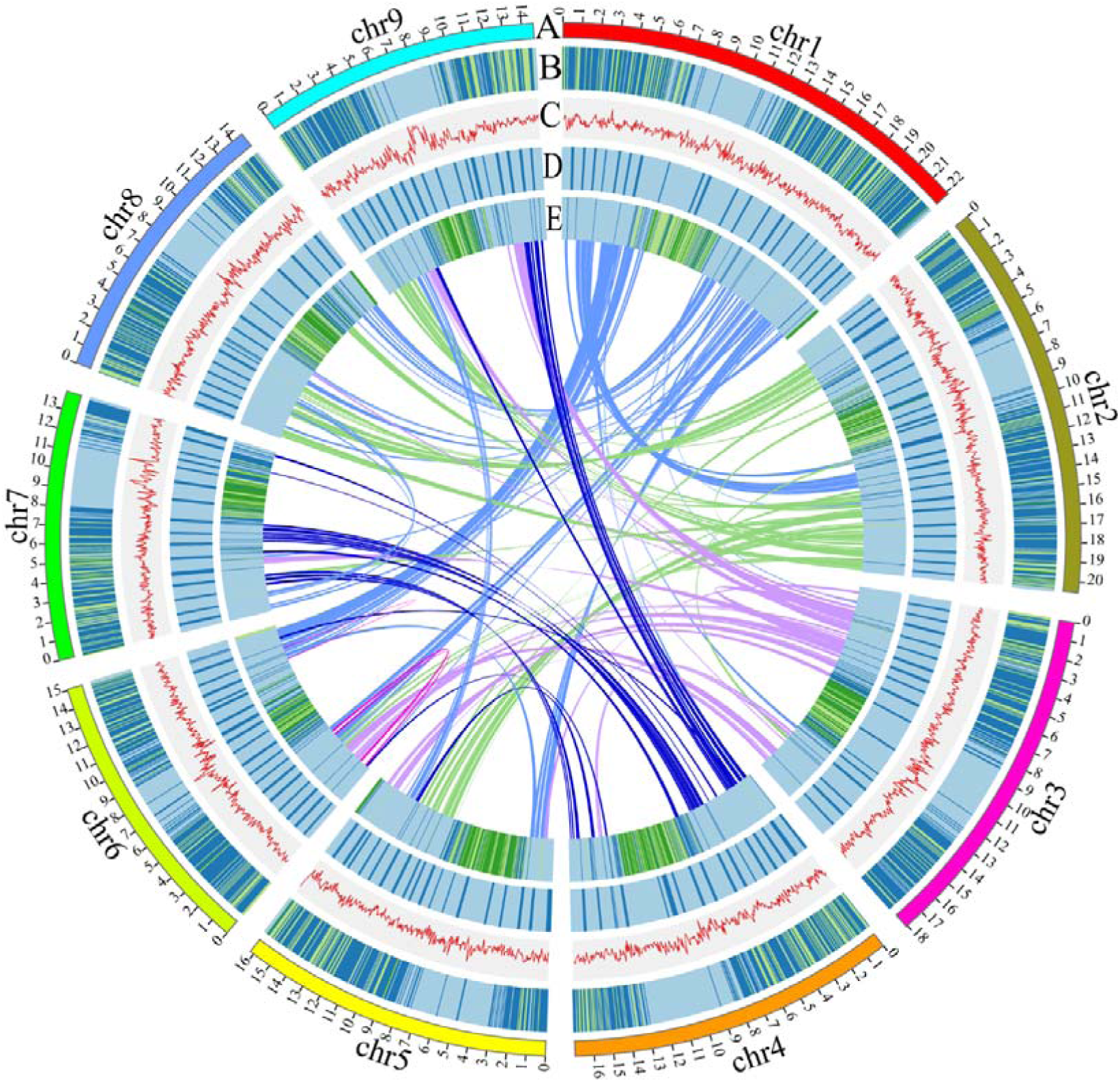
Genome landscape of *T. fournieri*. (A) The nine assembled chromosomes of *T. fournieri*. The distribution of (B) genes, (C) GC content, (D) SSRs, and (E) transposable elements (TEs). Darker colors correspond to higher gene density. Each linking line in the center of the Circos plot indicates a pair of homologous genes.

The combination of homology and *ab initio* gene prediction was used to label protein-coding genes in the *T. fournieri* genome, and 33532 genes were obtained (Table S6). The protein sequences produced by the predicted genes had an average length of 290 bp and an average of 6.48 exons per gene (Table S6). Using the Repeatmasker software, retroelements (10.9 Mb) accounted for 7.21% of the total sequence length (Table S7). In this study, we characterized the distribution of TEs and SSRs on the chromosomes of the *T. fournieri* genome. The results are presented in Fig. 1. To obtain the functional annotation information on the *T. fournieri* genome, we annotated all genes through the KEGG, NR, Swissprot, Tremble, KOG, GO, and Pfam databases (Table S8). 28812 genes were annotated through the Nr database (86.54%), 25095 genes through the GO database (75.37%), and 20162 genes through the KEGG database (60.56%) (Table S8), indicating a high degree of confidence in gene annotation.

### Comparative genomics analysis and evaluation

In order to study the genome evolution of the Linderniaceae family, where *T. fournieri* belongs, we studied and analyzed by comparative genomics four species of the Lamiaceae (*Salvia bowleyana*, *Origanum majorana*, *Scutellaria baicalensis* and *Ocimum basilicum*), two species of the Plantaginaceae (*Antirrhinum majus*, *Antirrhinum hispanicum*), one species of the Acanthaceae (*Strobilanthes cusia*), one species of the Phrymaceae (*Mimulus guttatus*), one species of the Amborellaceae (*Amborella trichopoda*), one species of the Vitaceae (*Vitis vinifera*), and two species of the Scrophulariaceae (*Lindenbergia philippehsis*) family, respectively, to a total of 12 species (Fig. 2A). The OrthoFinder software was used to obtain 34,150 homologous groups (Table S9), covering 424,454 genes (Table S10 and S11). Through the species evolutionary tree, *T. fournieri* was separated before the Plantaginaceae, Lamiaceae, Acanthaceae, and Scrophulariaceae during the Cretaceous period (103.38 Mya ago) (Fig. 2A). According to the gene family evolution calculations and analysis, 2423 gene families were expanded, and 3120 gene families were contracted (Fig. 2A, Table S12). Through Pfam annotation analysis on these expanded and contracted gene families, mainly gene families such as Hormone responsive protein, Ninja−family protein, Skp1 family, PA domain, LysM domain, C1 domain, and Transferase family were identified (Fig. S6). To estimate the polyploidy history of *T. fournieri*, we performed a curve-fitting analysis using the Ks distributions of the paralogs and orthologs identifed from *A. majus*, *S. baicalensis*, and *V. vinifera* (Fig. 2B). We observe that the Ks distribution of *T. fournieri* and *A. majus* has a main peak near 0.74, which was younger than the two peaks identified in the paralog analysis of *S. baicalensis* (0.93) and *V. vinifera* (1.32). In Fig. 2C and Fig. S7, there were a small number of collineated fragments in the dot plots of homologous genes of *T. fournieri*, *A. majus* and *V. vinifera*, and we speculate that a shared WGD event occurred in the common ancestor of *A. majus*, *V. vinifera* and *T. fournieri*.

**Fig. 2.**
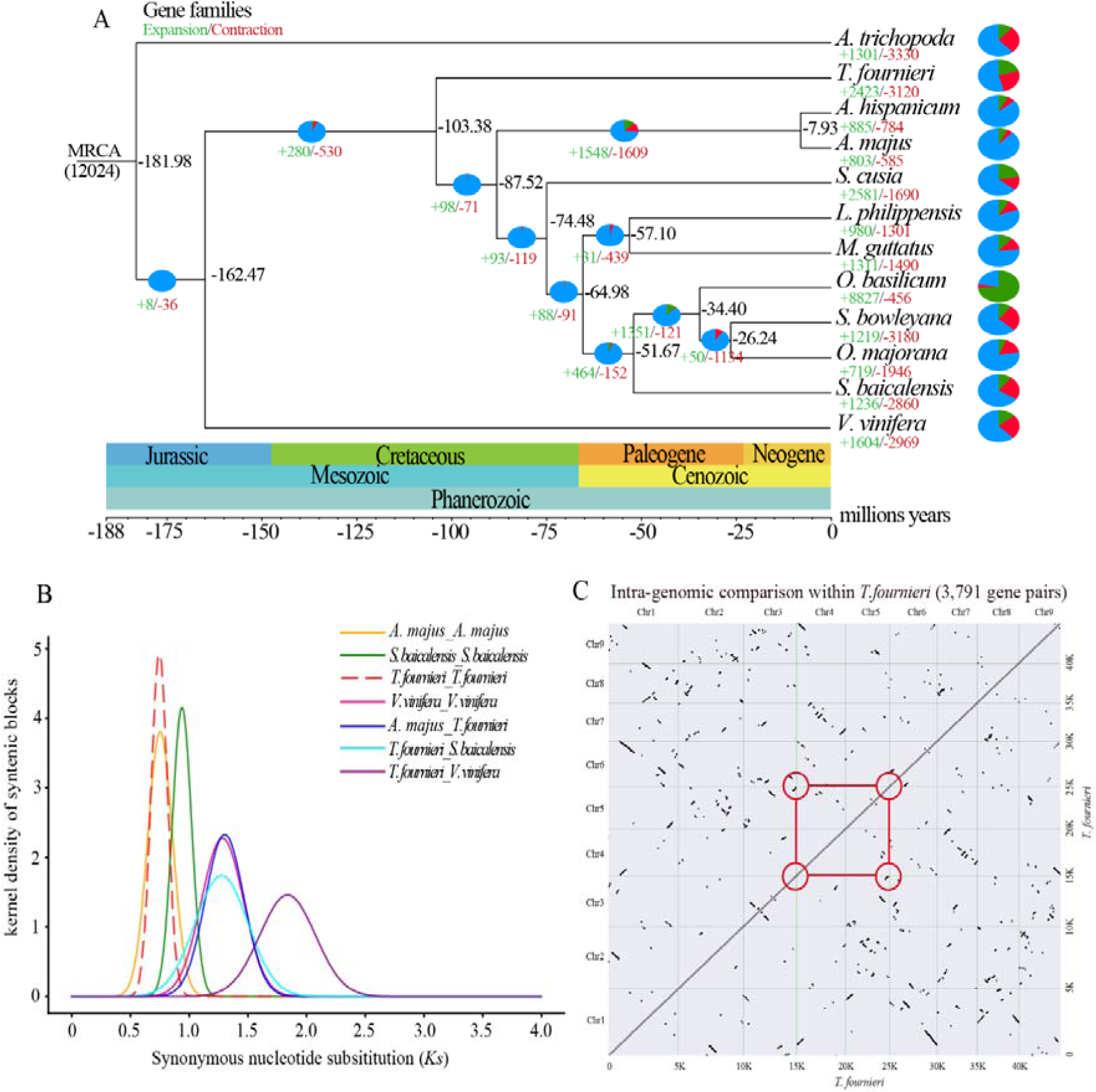
Comparative genomic and evolutionary analysis of *T. fournieri*. (A) Phylogenetic relationship between *T. fournieri* and 11 plant species. Green and red indicate the number of gene families that have expanded or contracted, respectively. The pie charts show the percentage of expanded (green), contracted (red), and conserved (blue) gene families among all gene families. Estimated divergence times (in millions of years) are shown in different colored sections below the phylogenetic tree. (B) The density distribution of homologous gene Ks values in *T. fournieri* and 11 plant species. (C) Dot plots of paralogs in the *T. fournieri* genome.

### Transcription and metabolism in *T. fournieri* flowers of different colors

The *T. fournieri* flowers are mainly composed of symmetrical petal lobes, a conical tube, and a flower neck. The flowers of different colors have in common that the flower neck is connected to the conical tube by the constriction area, both are yellow (Fig. S1), and all the mandibular petals have a macular patch (Fig. 3A). We sequenced the transcriptomes of these five differently colored *T. fournieri* flowers. The cleaned bases generated from each sample were about 6.5 G (15 samples sequenced in total), and the GC content was about 45% (Table S14). Through the Hisat software, the RNA-Seq data of the 15 samples were compared to the genome. The comparison efficiency was approximately 80%, indicating that the genome data and transcriptome data met the analysis requirements. 7308 Differently Expressed Genes (DEGs) were obtained using the DESeq2 software. By comparing White (W) with Deep blue (D), 2720 DEGs were obtained, among which 1136 DEGs were down-regulated, and 1584 DEGs were up-regulated (Fig. 3B). By comparing the White(W) with the Rose colored flowers, 1976 DEGs were obtained, among which 832 DEGs were down-regulated, and 1144 DEGs were up-regulated. By comparing White (W) with Blue and white (B), 1118 DEGs were obtained, with 510 DEGs down-regulated and 608 DEGs up-regulated. Comparing White(W) and Lemon drop(Y), 2431 DEGs were obtained, with 1149 DEGs down-regulated and 1282 DEGs up-regulated (Fig. 3B). Comparing W vs. Y, R, B, and D revealed a total of 155 DEGs, while 1177 specific DEGs were obtained when comparing W vs. Y (Fig. 3C). There were 1124 DEGs unique to the comparison of W vs. D. These DEGs were enriched in plant hormone signal transduction, anthocyanin biosynthesis, flavonoid biosynthesis, phenylalanine metabolism, and other metabolic pathways (Fig. S8A). There were 1177 DEGs unique to the W vs. Y comparison (Fig. 3C). These DEGs were mainly enriched in the MAPK signaling pathway, plant hormone signal transduction, flavonoid biosynthesis, phenylpropanoid biosynthesis, and other metabolic pathways (Fig. S8B). 199 DEGs were obtained by comparing the W vs. B flowers (Fig. 3C), enriched in metabolic pathways such as plant hormone signal transduction, flavonoid biosynthesis, and phenylpropanoid biosynthesis (Fig. S8C). 776 DEGs were obtained by comparing the W vs. R flowers (Fig. 3C) and were enriched in metabolic pathways such as plant hormone signal transduction, phenylpropanoid biosynthesis, and anthocyanin biosynthesis (Fig. S8D). These DEGs has affected a series of molecular response pathways in the plant, leading to different colors and morphological changes in the corolla of *T. fournieri*.

**Fig. 3.**
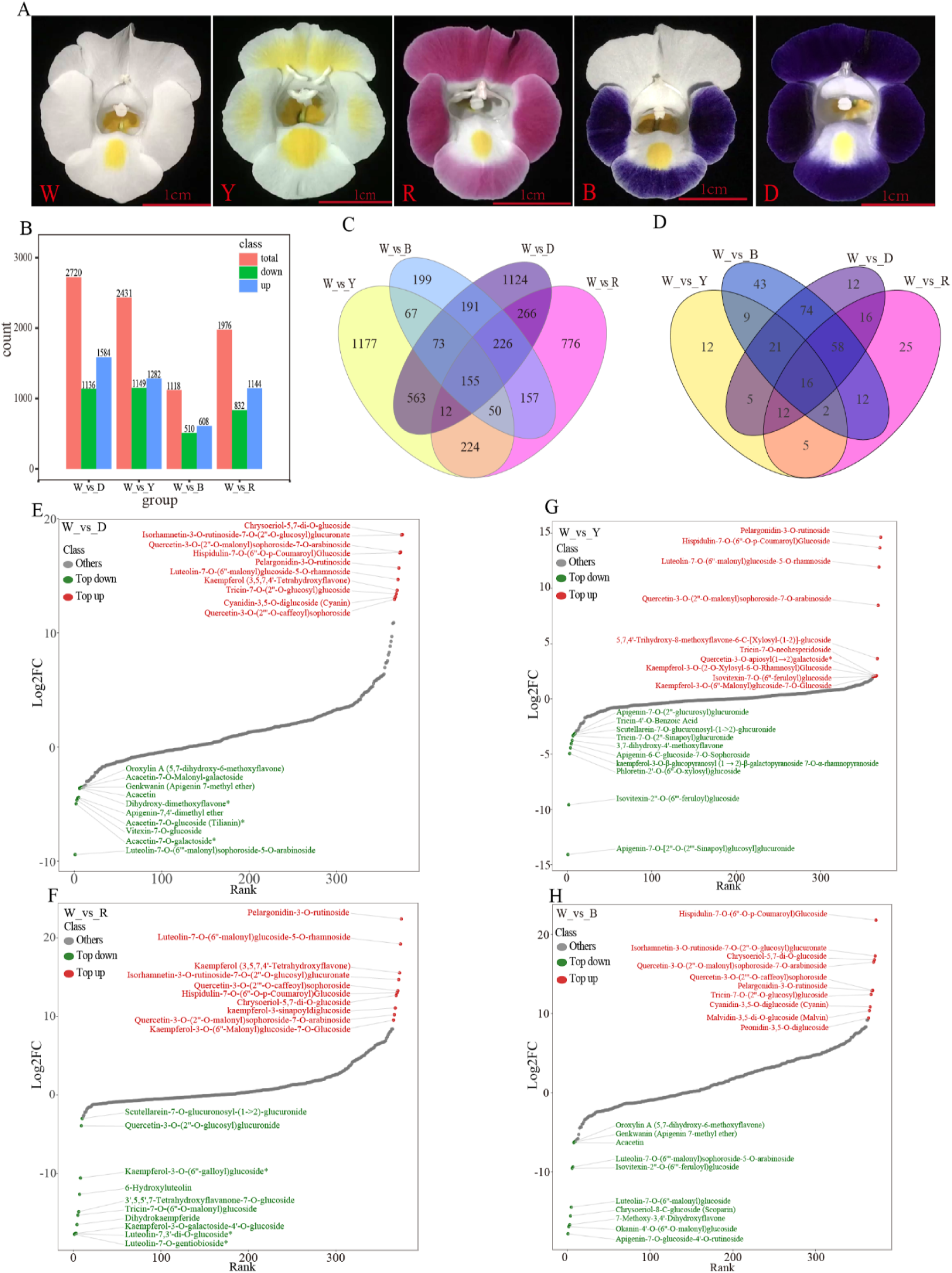
Transcriptome and metabolome results of *T. fournieri*. (A) Flowers of five different colors of *T. fournieri*. W, R, Y, B, and D represent white, rose, lemon drop, blue and white, and deep blue. (B) The number of up- and down-regulated genes between different groups was obtained by RNA-Seq analysis. (C) Venn diagram of differentially expressed genes between the different groups. Different colored dots represent different grouped samples. (D) Venn diagram of differentially accumulated metabolites between the different *T. fournieri* flower groups. The distribution of metabolite content differences in W_vs_D (E), W_vs_R (F), W_vs_Y (G), and W_vs_B (H) groups. Each dot represents an individual metabolite, green dots represent the top 10 down-regulated metabolites, and red dots represent the top 10 up-regulated metabolites.

We detected 375 flavonoid-related metabolites in the corolla of the five differentially colored *T. fournieri* flowers using UPLC-MS/MS (Table S15). By performing principal component analysis on the samples (including quality control samples), the results showed that the samples were almost clustered together. Indicating that the overall metabolite differences between the groups and the variability within the groups were small (Fig. S9A). By comparing Y with W, 82 significantly different metabolites were obtained, 44 were decreased, and 38 were increased in concentration. By comparing D with W, 214 significantly different metabolites were obtained, among which 52 decreased and 162 increased in concentration. By comparing R with W, 146 significantly different metabolites were obtained, of which 25 were decreased and 121 increased in concentration. By comparing B with W, 235 significantly different metabolites were obtained, among which 62 were decreased and 173 increased (Table S16). Finally, the comparison of Y, B, D, and R with W, revealed 12, 43, 12, and 25 unique metabolites with significant differences and 16 shared metabolites with significant differences (Fig. 3D, Table S16). Significantly different metabolites between the groups were obtained by orthogonal partial least squares discriminant analysis (OPLS-DA). Through the dynamic distribution diagram of the metabolite content differences between the W and D flower colors, we found that Quercetin-3-O-(2’’-O-malonyl)-sophoroside-7-O-arabinoside, Pelargonidin-3-O-rutinoside, and Cyanidin-3, 5-O-diglucoside (VIP>1) were significantly increased (Fig. 3E, Fig. S9B). When comparing R with W, Pelargonidin-3-O-rutinoside (VIP>1) and Luteolin-7-O-(6’’-malonyl)-glucoside-5-O-rhamnoside were significantly increased in concentration (Fig. 3F, Fig. S9E). Comparing W with Y, Pelargonidin-3-O-rutinoside (VIP>1), Quercetin-3-O-(2’’-O-malonyl)-sophoroside-7-O-arabinoside and Quercetin-3-O-apiosyl (1 →2)-galactoside was signific-antly accumulated in W (Fig. 3G, Fig. S9C). Similarly, when B was compared with W, we found that Isorhamnetin-3-O-rutinoside-7-O-(2’’-O-glucosyl)-glucuronate, Pelargonidin-3-O-rutinoside, Cyanidin-3,5-O-diglucoside, Malvidin -3,5-di-O-glucoside and Peonidin-3,5-O-diglucoside were significantly increased in concentration (Fig. 3H, Fig. S9D).

We detected a total of 108 anthocyanin-related metabolites in the five differently colored *T. fournieri* corollas by using UPLC-MS/MS, of which 58 anthocyanins were detected (Table S17). We performed UV (unit variance scaling) processing on those 58 metabolites and drew a cluster heat map. Two additional Pelargonidin metabolites were identified in sample W, while sample R contained multiple Pelargonidin and Peonidin related metabolites, and samples B and D mainly contained Malvidin, Cyanidin, and Peonidin related metabolites (Fig. 4A). Through a metabolite content histogram, we found that sample B contained a high concentration of Cyanidin-3-O-(6-O-p-coumaroyl)-glucoside (Fig. S11), and sample D contained a high concentration of Cyanidin-3-O-(6-O-malonyl-beta-D-glucoside), Cyanidin-3-O-glucoside and Cyanidin-3,5-O-diglucoside (Fig. 4A, Fig. S11). However, samples R, B, and D contained a high concentration of Delphinidin-3-O-(6-O-p-coumaroyl)-glucoside and Delphinidin-3,5-O-diglucoside relative to W, Y (Fig. 4A, Fig. S12). In terms of peonidin metabolites, we found that samples B and D contained a high concentration of Peonidin-3-O-(6-O-p-coumaroyl)-glucoside, Peonidin-3,5-O-diglucoside and Peonidin-3-O-glucoside (Fig. 4A, Fig. S13). Similarly, samples B and D also contained small amounts of petunidin-related metabolites, such as Petunidin-3-O-glucoside, Petunidin-3-O-galactoside, Petunidin-3-O-sambubioside-5-O-glucoside, and Petunidin -3-O-(6-O-malonyl-beta-D-glucoside) (Fig. 4A, Fig. S14). In the rose-colored R sample, a large number of pelargonidin metabolites were identified, such as Pelargonidin-3-O-rutinoside, Pelargonidin-3-O-glucoside, Pelargonidin-3,5-O-diglucoside, Pelargonidin-3-O-galactoside, Pelargonidin -3-O-sambubioside, Pelargonidin-3-O-(6-O-malonyl-beta-D-glucoside) and Pelargonidin-3-O-sophoroside (Fig. 4A, Fig. S15). There were numerous mallow pigment-related metabolites in the blue-colored B and D samples, such as Malvidin-3-O-sambubioside-5-O-glucoside, Malvidin-3,5-O-diglucoside, Malvidin-3-O-(6-O-p-coumaroyl) -glucoside and Malvidin-3,5-O-diglucoside (Fig. 4A, Fig. S16). In terms of flavonoid metabolites in different samples, sample B contained a high concentration of Rutin (Fig. 4A, Fig. S17) and Kaempferol-3-O-rutinoside (Fig. 3G, Fig. S19 C). Sample R contained a high concentration of Dihydrokaempferol (Fig. 3G, Fig. 19B), Naringenin (Fig. 3G, Fig. S19D), and Naringenin-7-O-glucoside (Fig. 3G, Fig. S19E). The above results indicated that different concentrations of anthocyanin glycosides were the main contributors to the different coloration of *T. fournieri* flowers.

**Fig. 4.**
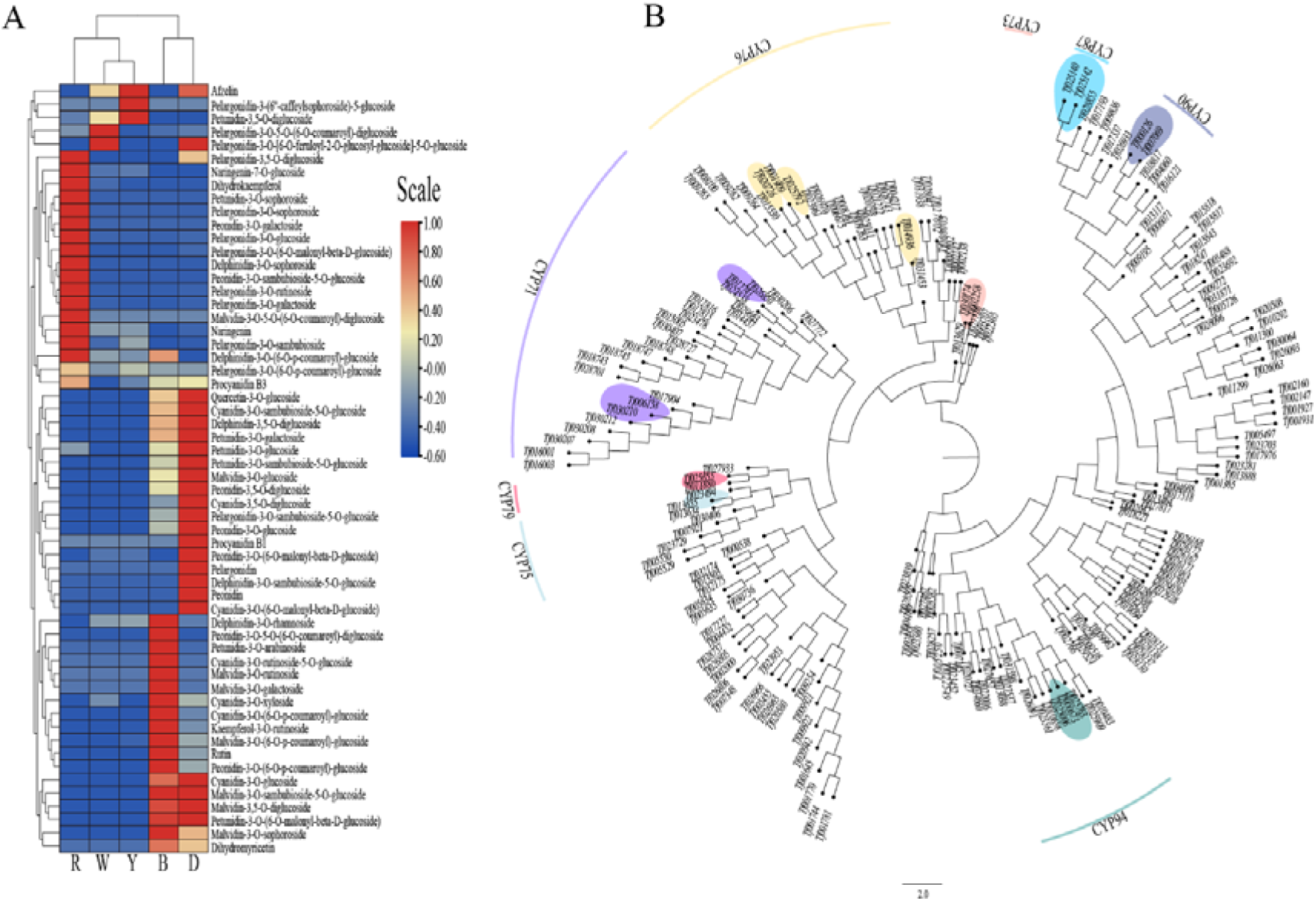
Analysis of anthocyanin content and the cytochrome P450 gene family in *T. fournieri*. (A) Heatmap of anthocyanin metabolite content between different sample groups. The anthocyanin metabolite data were processed by UV (unit variance scaling). Cluster heatmaps were drawn using the R program heatmap package. (B) Evolutionary tree of genes encoding cytochrome P450 proteins in *T. fournieri*. The outermost circle of the phylogenetic tree indicates the subfamilies corresponding to the different tree branches. The genes highlighted with different colors correspond to DEGs. The cytochrome P450 family members were clustered by neighbor ligation using the iqtree software (-bb:1000; MFP).

### Phylogenetic analysis of gene families

The plant cytochrome (CYP) P450 gene family plays important regulatory and catalytic roles in plant growth, development, and secondary metabolite biosynthesis(Hansen *et al*., 2021). In this study, we downloaded the protein sequences from all the members of the *A. thaliana* cytochrome P450 gene family and used blastp to make a global alignment with the corresponding *T. fournieri* protein sequences. Through sequence alignment analysis, we initially obtained 216 cytochrome P450 protein sequences, and a total of 193 P450 protein sequences were obtained based on the annotation information and the removal of redundant sequences. According to the classification based on the *A. thaliana* subfamily information, the P450 gene family can be divided into 39 subfamilies (Table S18). The developmental tree of the P450 gene family was constructed using the iQtree software. The CYP71 subfamily had the largest subfamily branch, containing 23 genes (Fig. 4B). According to the gene FPKM values from the transcriptome of the differently colored flowers, we obtained the expression information of all P450 gene families in *T. fournieri* (Fig. S18). By using the |log2Fold Change| >= 1 cutoff, we screened genes with significant expression differences, which have been marked with different colors in Fig. 4B. Similarly, the multimember Myb gene family plays important roles in regulating plant growth, development, and anthocyanin biosynthesis(Yang *et al*., 2022). In this study, we downloaded the *A. thaliana* R2R3-Myb gene family information and obtained 62 *T. fournieri* R2R3-Myb genes through sequence alignment (Table S19). Using the *A. thaliana* and *T. fournieri* Myb protein sequences, the Myb transcription factor family phylogenetic tree was constructed (Fig. S19). The S32 branch was the largest subfamily branch with 16 genes. Genes in different subfamilies were expressed differently in the flowers with different colors, and they might play an important role in regulating anthocyanin biosynthesis, which is responsible for the formation of different flower colors.

### Biosynthetic pathways of flavonoids and anthocyanins in *T. fournieri*

Flavonoids and anthocyanins play key roles in plant growth, development, and organ coloration(Zhao *et al*., 2022). In this study, we identified all the key genes involved in the flavonoid synthesis pathway to reveal their functions in the formation of different flower colors. We identified 37 *4CLs*, 2 *ANRs*, 1 *ANS*, 13 *CHIs*, 6 *CHSs*, 3 *CYP73As*, 6 *DFRs*, 1 *F3’5’H*, 7 *UFGTs*, 4 *F3Hs*, 4 *F3’Hs*, 4 *FLSs*, 8 *HCTs*, 2 *LARs* and 6 *PALs* (Table S20). We screened the genes in the pathway with significant differences and drew a flavonoid regulatory network map (Fig. 5). By comparing the FPKM values of different genes involved in flower color formation (Fig. 5), we clearly found that the ANR enzyme gene *Tf014160* was differentially expressed in each group, with low expression in R and D samples and high expression in W, Y and B samples.

**Fig. 5.**
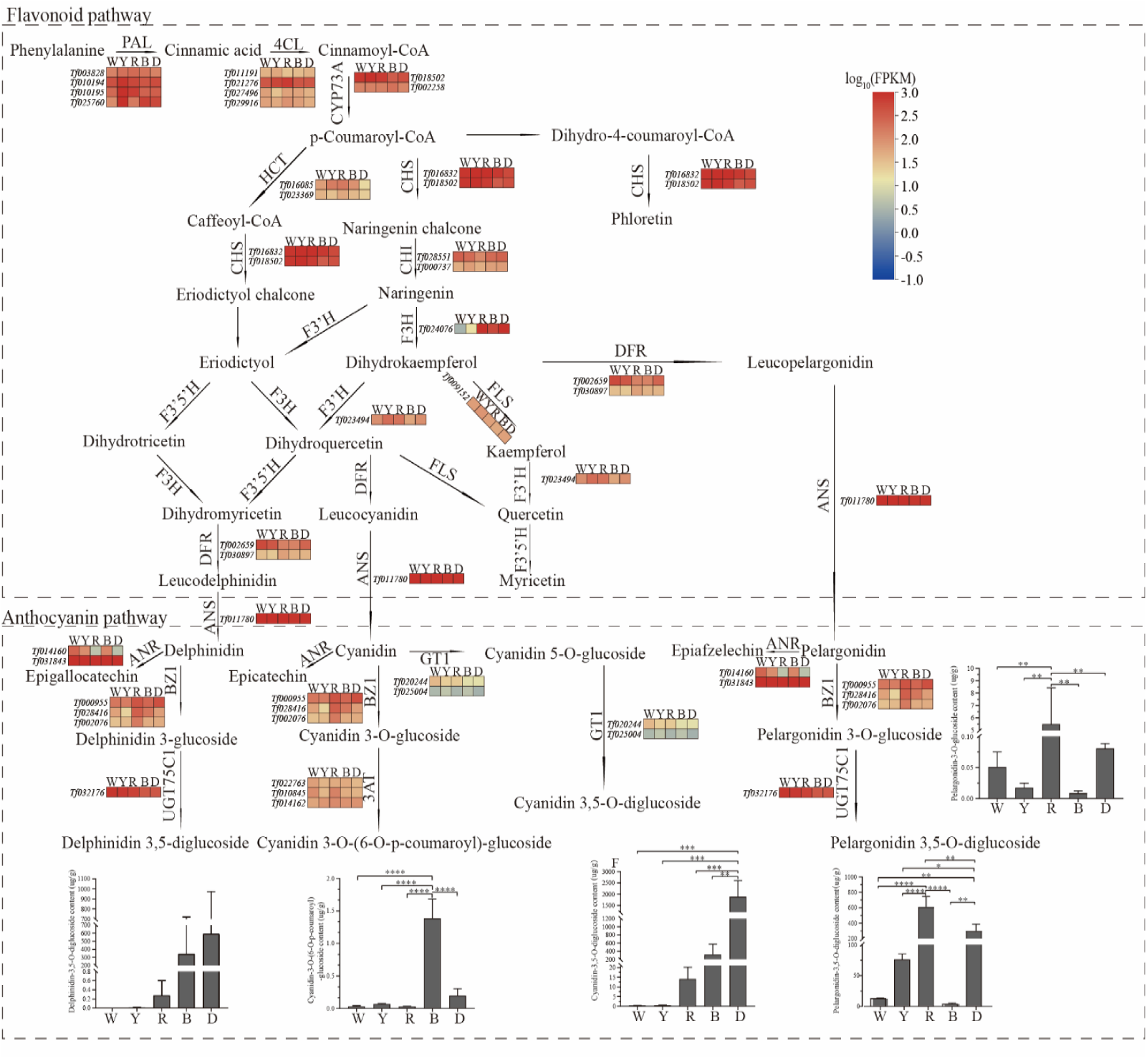
Key pathways for flavonoid accumulation and anthocyanin synthesis in *T. fournieri*. Heatmap of flavonoid accumulation and expression levels of candidate genes involved in anthocyanin synthesis in different flower color tissues of *T. fournieri*. Red and blue correspond to high and low expression levels of the related genes in the pathway, respectively(log_10_(FPKM)). Histograms illustrate the content of related anthocyanin metabolites in the different flower color tissues of *T. fournieri*, prepared by GraphPad Prism 8. (*<0.05, **<0.01, ***<0.001, ****<0.0001)

The *F3H* is one of the key enzymes in the flavonoid metabolic pathway, significantly impacting the biosynthesis of flavonoids(Li *et al*., 2020). The combined transcriptome and metabolome analysis revealed that the F3H enzyme gene *Tf024076* was significantly differentially expressed between the groups (Fig. S20). The FPKM value of *Tf024076* in R was 2866.32, 974.94-fold higher compared to that of W, and 184.92-fold higher compared to that of Y. We speculate that the F3H enzyme gene *Tf024076* plays an important role in regulating the coloration of the *T. fournieri* flower. Anthocyanin synthase (ANS) is the most critical enzyme in the process of anthocyanin synthesis and transformation(Sharma *et al*., 2022). In this study, we found that the FPKM values of the ANS enzyme *Tf011780* gene in the W, Y, and R samples were 1961.70, 2709.95, and 1834.65, and in the B and D samples were 883.65 and 894.48. *Tf011780* positively regulates the anthocyanin biosynthesis pathway of *T. fournieri*. In this study, we identified the genes involved in the anthocyanin synthesis pathway and found 11 *3ATs*, 16 *BZ1s*, 1 *FG3*, 2 *GT1s*, 8 *HIDHs*, 14 *PTSs*, and 1 *UGT75C1* (Table S21). In this study, we screened 2 *GT1s*, and we speculated that the differential expression of this gene in each group affected Cyanidin-3,5-O-diglucoside synthesis. The Cyanidin-3,5-O-diglucoside content in each sample was significantly different. W, Y, and R samples had a significantly lower content than B and D (*P*<0.001). Similarly, the differential expression of *BZ1* and *UGT75C1* genes across the flower color types eventually led to significant differences in the contents of Delphinidin-3,5-O-diglucoside, Cyanidin-3-O-(6-O-p-coumaroyl)-glucoside, Pelargonidin-3,5-O-diglucoside, and Pelargonidin-3-O-glucoside. Due to their significant differential expression, we speculate that these genes have a positive role in the anthocyanin glycoside synthesis pathway.

Through the joint transcriptome and metabolome analysis, we evaluated the genes involved in the synthesis of anthocyanin glycosides. Firstly, correlation analysis was performed on the quantitative values of all samples in each groups. Data with a correlation coefficient greater than 0.85 and a p-value less than 0.5 were selected (Table S22). The screened data were used to draw a correlation clustering heat map (Fig. S20) using the Pretty Heatmaps package of R. We found that the *Tf024076* was positively correlated with Pelargonidin, and Pelargonidin-3-O-glucoside was significantly positively correlated with Tf024076 (*P*<0.0001) (Table S22). Therefore, *Tf024076* potentially plays an active role in the synthesis of Pelargonidin-related metabolites. Based on the DEGs that are part of the anthocyanin synthesis pathway, we screened 23 related genes and measured the expression fold change by real-time quantitative PCR (RT-qPCR). We compared the log2(RT-qPCR) values of these 23 genes with the RNA-Seq data, which showed that the expression of these selected genes in our transcriptome dataset was highly consistent with the qRT-PCR results (Fig. S23).

## Discussion

*T. fournieri* is an ornamental plant with high economic value. It has many flowers and rich colors and is a model plant for studying angiosperm fertilization and development (Kikuchi *et al*., 2006; Liu *et al*., 2020). In this study, we reported the generation of a 164.4Mb *T. fournieri* genome (Table S4.3), including its abundant repetitive elements and annotated information (Table S6). Our phylogenetic analysis clearly reveals the early evolutionary relationships of *T. fournieri*. Specifically, its relationship with Lamiaceae (*S. bowleyana*, *O. majorana*, et al.), Plantaginaceae (*A. majus*, *A. hispanicum*), Acanthaceae (*S. cusia*), Scrophulariaceae (*L. philippehsis*), Phrymaceae (*M. guttatus*), Amborellaceae (*A. trichopoda*), and Vitaceae (*V. vinifera*) (Fig. 2A). Through the phylogenetic l tree analysis, it was found that *T. fournieri* diverged earlier than Lamiaceae and Plantaginaceae. By comparing the *T. fournieri* chloroplast genomes, it was also found that *T. fournieri* was significantly differentiated from Plantaginaceae, Acanthaceae, and Scrophulariaceae(Chen *et al*., 2021). According to the evolutionary relationship analyses, *T. fournieri* belongs to the genus *Torenia* of the family Linderniaceae rather than the genus *Torenia* of the family Scrophulariaceae, as currently listed in the Angiosperm Phylogeny Group (APG) III classification system. By combining Ks and phylogenetic analysis, we concluded that the WGD events in *T. fournieri* occurred at the same time as that of *A. majus* (Fig. 2C, Fig. S7A). Therefore, our assembled *T. fournieri* genome provides new insights and resources for the comparative study of Linderniaceae.

In this study, we analyzed the differences underlying the five different flower color types of *T. fournieri* japonica through transcriptomics and metabonomics. RNA-Seq analysis revealed that many DEGs were differentially expressed in samples of different flower colors. These DEGs were involved in various metabolic pathways (such as flavonoid and anthocyanin pathways) (Fig. S8). Plant cytochrome P450 genes and R2R3-MYB transcription factors participate in various biochemical pathways and produce a variety of metabolites (such as phenylpropanoids, terpenes, cyanogenic glycosides, and glucosinolates, etc.), which play an important role in flavonoid biosynthesis and their colored compounds anthocyanins(Distefano *et al*., 2021; Ma and Constabel, 2019; Nguyen and Dang, 2021). The *F3’H* and *F3’5’H* genes belong to the CYP75 gene subfamily(Tanaka and Brugliera, 2013). The inhibition of the *F3’5’H* gene leads to the increase of anthocyanins(He *et al*., 2013; Tanaka and Brugliera, 2013). In contrast, the increased expression of the *F3’H* gene in *T. fournieri* increases the anthocyanin content and leads to a pink flower color(Tanaka and Brugliera, 2013). We thoroughly assessed the P450 gene family by analyzing the *T. fournieri* genome and identified a total of 39 P450 subfamilies (Fig. S18, Table S18), among which the CYP75 subfamily contains eight genes. These genes include 4 *F3’H* genes (Table S20), which were differentially expressed in different samples and potentially positively regulate the formation of color in *T. fournieri* flowers. We found that the CYP87 subfamily *Tf028855* gene was significantly highly expressed in the R flowers compared to the other flower types (Table S18). Moreover, the content of Pelargonidin-3-O-rutinoside in the R flowers was also significantly higher than that in W, Y, B, and D flower types (Fig. S15A). A joint analysis revealed that the *Tf028855* gene was significantly positively correlated with Pelargonidin-3-O-rutinosid accumulation (*P*<0.001). Therefore, Tf028855 is potentially a key gene that regulates the synthesis pathway of Pelargonidin-3-O-rutinosid, resulting in the rose color in R flowers. Several genes belonging to these gene families are differentially expressed in the different flower types, directly or indirectly affecting the metabolism of *T. fournieri* flavonoids or anthocyanins, playing an important regulatory role in *T. fournieri* corolla coloring.

The purple and blue flowers mainly contain anthocyanidins, delphinidin, and its methylated derivatives, petunidin, and malvidin(Mekapogu *et al*., 2020). The anthocyanins of the magenta, red, scarlet, and pink-colored flowers are mainly Pelargonidin, Cyanidin, Delphinidin, Peonidin, Petunidin, and Malvidin(Fu *et al*., 2021; Iwashina, 2015). Similarly, our study found no significant difference in cyanidin content in the rose-colored R flower types (Fig. S11). However, the content of Delphinidin, Pelargonidin, Peonidin, Petunidin, Malvidin, Naringenin, and Dihydrokaempferol-related glycosides was significantly higher than that of anthocyanins in other samples (Fig. 4A, Fig. S12A, Fig. S15A, Fig. S15B, Fig. S15G -K, Fig. S17B, Fig. S17D, and E; Table S17). We found that the B and D flower types mainly contained Quercetin, Cyanidin, Delphinidin, petunidin, malvidin, Pelargonidin, and Peonidin related glycosides. We also found that the Rutin and Kaempferol-3-O-rutinoside contents in the B-colored flowers were also significantly higher than those in the other flower colors (Fig. S17A and Fig. S17C). In the yellow-colored Y flowers, we found that they mainly contained Afzelin, Pelargonidin-3-O-(6’’-ferulylsambubioside)-5-O-(malonyl)-glucoside, and Petunidin-3,5-O-diglucoside (Fig. 4A, Table S17). The differences in anthocyanin metabolites indirectly revealed the mechanisms underlying the different flower colors of *T. fournieri*.

Anthocyanin glycoside is an important water-soluble flavonoid compound in plants, mainly present in tissues such as flowers, fruits, and leaves, resulting in the different color appearance of these tissues (Mizuno *et al*., 2021; Park *et al*., 2018). The MBW complex (R2R3-MYB, bHLH, and WD40) proteins play an important regulatory role in the plant anthocyanin synthesis regulatory pathway, and the R2R3-MYB transcription factors are critical regulators of plant anthocyanin synthesis(Li, 2014; Ma and Constabel, 2019). In the Petunia R2R3-MYB gene family, anthocyanin synthesis regulators (ASR) can participate in the WMBW (WRKY, MYB, B-HLH, and WDR) anthocyanin regulatory complex by interacting with the AN1 and AN11 transcription factors, thus regulating the different flower color formation in Petunia(Zhang *et al*., 2019). Similarly, the *NnMYB5* transcription factor of the lotus R2R3-MYB gene family is a transcriptional activator of anthocyanin synthesis, playing an important role in regulating flower color(Sun *et al*., 2016). Therefore, in this study, we identified the R2R3-MYB transcription factors present in the *T. fournieri* genome and obtained 62 related genes, of which 11 belong to the S32 R2R3-MYB subfamily (Fig. S19, Table S19). Among these transcription factors, we found that *Tf023331* had the highest expression in the D flower type, which also exhibited a significantly higher content of Malvidin-3-O-glucoside compared to other samples (Fig. S16E). Thus, *Tf023331* may be a key gene involved in the Malvidin-3-O-glucoside synthesis pathway, and Malvidin-3-O-glucoside may be responsible for the darker color of the D flowers compared with the other flower colors.

Anthocyanin glycoside synthesis is mainly catalyzed by a series of enzymes and transported to the vacuole for storage through various modifications(Tanaka *et al*., 2008). These enzymes mainly include CHS, CHI, F3H, F3’H, FLS, FNS, DFR, ANS, ANR, and GT1. In the *T. fournieri* genome, we identified 37 4*CLs*, 2 *ANRs*, 1 *ANS*, 13 *CHIs*, 6 *CHSs*, 3 *CYP73As*, 6 *DFRs*, 1 *F3’5’H*, 7 *UFGTs*, 4 *F3Hs*, 5 *F3’Hs*, 4 *FLSs*, 8 *HCTs*, 2 *LARs*, 6 *PALs*, 11 3*ATs*, 16 *BZ1s*, 1 *FG3*, 2 *GT1s*, 8 *HIDHs*, 14 *PTSs* and 1 *UGT75C1* genes (Table S20 and S21). ANS was shown to be a key enzyme in the anthocyanin biosynthesis pathway, catalyzing the conversion of colorless anthocyanins into colored anthocyanins(Sharma *et al*., 2022). *SmANS* is an anthocyanin synthase gene in the downstream anthocyanin biosynthesis pathway. Low expression of the *SmANS* leads to the production of white flowers from purple flowers in *Salvia miltiorrhiza(Lin et al., 2022)*. *SmANS* overexpression of promoted anthocyanin accumulation in *S. miltiorrhiza* and restored the purple flower phenotype(Li *et al*., 2019). In the *Dendrobium officinale* anthocyanin biosynthesis pathway, *DoANS* and *DoUFGT* encoding an anthocyanin synthase and a UDP-glucose flavonoid-3-O-glucosyltransferase, respectively, are key regulatory genes associated with anthocyanin differential accumulation(Yu *et al*., 2018). We combined the metabolome and transcriptome data analysis and found that *TfANS* may be the key gene that determines the color of the perianth segments of *T. fournieri*. Flavanone 3-hydroxylase (F3H) plays an important role in the flavonoid biosynthetic pathway. The expression of *TfF3H* in a white *T. fournieri* perianth is lower than that in a purple perianth. When *TORE1* (Torenia retrotransposon 1) was inserted into the 5’ upstream region of the *TfF3H* gene in white perianth flowers, it activated the expression of *TfF3H*, which resulted in the white flowers turning pink(Nishihara *et al*., 2014). In this study, we found that *Tf024076* was expressed very lowly in the white W and yellow Y flower types, but was highly expressed in the rose R, blue B and D flower types. We also combined the *Tf024076* transcript expression information with the metabolome data, and observed that *Tf024076*, 3,4,2’,4’,6’-Pentahydroxychalcone-4’-O-glucoside, Naringenin-7-O-glucoside and Pelargonidin-3-O-glucoside were significantly positively correlated (Fig. S20). However, the Naringenin-7-O-glucoside and Pelargonidin-3-O-glucoside contents in the R flower types were significantly higher compared to the other flower types (Fig. 4A, Fig. 15B, Fig. S17E). Therefore, *Tf024076* has a positive regulatory effect in the biosynthesis of Naringenin-7-O-glucoside and Pelargonidin-3-O-glucoside, and these two anthocyanin metabolites are also responsible for the rose color of the R flower type.

In conclusion, the assembled *T. fournieri* sequence provides a reference genome for the Linderniaceae, serving as a valuable resource for future genome editing research and molecular marker-assisted breeding. It also provides insights into the evolution of the genus Torenia in the Linderniaceae. The RNA-Seq data from flowers of different colors of *T. fournieri* revealed differences at the molecular level, and the metabolome data revealed differences at the biochemical level. The integrated analysis shed light on the mechanisms underlying the corolla colors in *T. fournieri*. Importantly, the genes and metabolites identified in this study further provide a multi-omics resource for understanding the growth, coloration, and antioxidant properties of *T. fournieri* corolla.

## Acknowledgements

We acknowledge Drs. Shuo Li, Cai Gao and Zhongxing Li for their help and advice during the experiment. We gratefully thank Professor Qian Li (Xinjiang Agriculture University) for the helpful advice and discussion of this manuscript.

## Funding

This study was funded by China Agriculture Research System of MOF and MARA(CARS-34) and the Wetland and Grassland Research Center of Shaanxi Academyof Forestry(SXLK-ZX-2021-06).

## Author Contributions

TH and PY planted and designed the study; JS analysed data and wrote the manuscript; JJ, SL and HK data collection and performed experiments; JY, NM and RY analyzed data and planned the experiments;YC and YW edited and revised the manuscript.

## Declarations

The authors declare that they have no conflicts of interest associated with this work.

## Data Availability Statement

The transcriptome and genome sequencing data of *T. fournieri* have been deposited in NCBI under the bioproject Accession PRJNA928569 and PRJNA928860.

## Notes

### Competing Interest Statement

The authors have declared no competing interest.

